# High School Science Fair: School Location Trends in Student Participation and Experience

**DOI:** 10.1101/2023.02.01.526679

**Authors:** Frederick Grinnell, Simon Dalley, Joan Reisch

## Abstract

The findings reported in this paper are based on surveys of U.S. high school students who registered and managed their science and engineering fair (SEF) projects through the online Scienteer website over the three years 2019/20, 2020/21, and 2021/22. Almost 2500 students completed surveys after finishing all their SEF competitions. We added a new question in 2019/20 to our on-going surveys asking the students whether their high school location was urban, suburban, or rural. We learned that overall, 74% of students participating in SEFs indicated that they were from suburban schools. Unexpectedly, very few SEF participants, less than 4%, indicated that they were from rural schools, even though national data show that more than 20% of high school students attend rural schools. Consistent with previous findings, Asian and Hispanic students indicated more successful SEF outcomes than Black and White students. However, whereas Asian students had the highest percentage of SEF participants from suburban vs. urban schools – 81% vs. 18%, Hispanic students had the most balanced representation of participants from suburban vs. urban schools – 55% vs. 39%. Differences in students’ SEF experiences based on gender and ethnicity showed the same patterns regardless of school location. In the few items where we observed statistically significant (probability <.05) differences based on school location, students from suburban schools were marginally favored by only a few percentage points compared to students from urban schools. In conclusion, based on our surveys results most students participating in SEFs come from suburban schools, but students participating in SEFs and coming from urban schools have equivalent SEF experiences, and very few students participating in SEFs come from rural schools.

## Introduction

In recent years, ideas about how best to accomplish science education have focused increasingly on hands-on science and engineering (S&E) practices [1–3]. Science and engineering fairs (SEFs) offer one way that students can experience for themselves the practices of science and engineering described by Next Generation Science Standards (NGSS) [4, 5].

SEFs potentially can promote three important and desirable outcomes: (i) mastery of S&E practices; (ii) interest in science; and (iii) interest in science careers [6]. The idea that SEF participation can have a positive impact on high school students is consistent with research showing that science project-based learning advances students’ STEM understanding and interests at both the high school [7–9] and undergraduate levels [10–12].

The number of U.S. high school students who participate in SEFs each year is not known exactly. However, the 2009/12 NCE-HSLS report found that 5% of high school students participated in science competitions during high school [13], which would correspond to an upper limit of about 750,000 students given the overall U.S. public high school population of about 15 million [14] (does not include about 10% of students attending private schools [15]). Similarly, about 5% of the almost 16,000 undergraduate students surveyed in the college Outreach Programs and Science Career Intentions survey (students in introductory freshman, mostly English, classes) reported participating in SEFs in high school [16].

A national study of middle schools identified three major SEF types: mandatory SEFs with high support (curriculum, class time, teacher engagement) (23% of students); mandatory with low support (57% of students); and voluntary with low support (20% of students) [6]. Teacher support for students was more limited in high poverty schools and schools with a higher proportion of Black students [17]. Higher income parents and those parents with greater educational attainment were more likely to provide greater SEF support for their children [18].

Some national studies with high school students suggested that the main overall effect of participating in SEFs was retention of students who already were interested in S&E [16, 19]. Consistent with this possibility, we found that students who participated in SEFs in 11^th^ and 12^th^ grades were more likely to be interested in careers in S&E compared to students in 9^th^ and 10^th^ grades and more likely to indicate that SEF participation increased their interests [20]. However, most high school students who participated in SEFs were required to do so, and that requirement decreased the positive impact of SEFs on the students [21]. Also, the national cohort of students in our studies indicated that help from scientists and teachers was more important than help from parents, and the most common source of help for students was the internet [22].

Innovative high school programs that combined student participation in SEFs with student and teacher support promoted STEM engagement and learning for all students including those from under-represented ethnic minorities and low socioeconomic backgrounds [23–27]. We found that experiential factors such as help from scientists, coaching for the SEF interview and help fine-tuning the SEF report all correlated with greater likelihood that students indicated SEF participation increased their S&E interests [22]. Earlier work by others also showed that access to outside of school facilities [28] and research resources [29] enhanced students’ SEF experiences.

Because the overall goal of our research has been to establish a base of knowledge regarding student experiences in high school SEFs, we periodically add new questions to our ongoing surveys. For the cohort of high school students that we surveyed beginning in 2018/19, we added a new question about ethnicity. Ethnicity trends with students in the 2018/19 and 2019/20 survey groups showed two important sets of ethnicity-dependent differences. First, compared to the general high school student population, Asian students were over-represented in SEFs by 5-fold or more, whereas students in other ethnic groups were under-represented. Overall, the ethnic distribution of student SEF participants was similar to the percentages of students in the national NCE-HSLS (2009) who indicated that they planned to pursue a STEM major when they reached college [30]. Second, Asian and Hispanic students had more positive SEF outcomes than Black and White students [22], a pattern different from general studies of ethnicity in relationship to student academic achievement [31, 32].

In 2019/20, we added another new question in our surveys. this one asking students whether the location of their high school was urban, suburban, or rural. In this paper, we report findings for this question with the students who completed SEF surveys in 2019/20, 2020/21, and 2021/22. According to the survey results, most students participating in SEFs came from suburban schools, but students participating in SEFs and coming from urban schools had equivalent SEF experiences. Unexpectedly, very few students participating in SEFs came from rural schools. Details are reported herein.

## Materials and methods

This study was approved by the UT Southwestern Medical Center IRB (#STU 072014-076). Study design entailed administering to students a voluntary and anonymous online survey using the REDCap survey and data management tool [33]. Survey recipients were U.S. high school students who participated in SEFs during the 2019/20, 2020/21, and 2021/22 school years using Scienteer (www.scienteer.com) for online SEF registration, parental consent, and project management. Although we treat the Scienteer SEF population as a national group of U.S. high school students, it should be recognized that these students come from only 7 U.S. states: Alabama, Louisiana, Maine, Missouri, Texas, Vermont, and Virginia. We have no information about the locations where SEF fairs are held in the seven states.

After giving consent for their students to participate in SEFs, parents could consent for their students to take part in the SEF survey. However, to prevent any misunderstanding by parents or students about a possible impact of agreeing to participate or actually participating in the survey, access to the surveys was not available to students until after they finished <u>all</u> of their SEF competitions. When they initially registered for SEFs, students whose parents gave permission were told to log back in to Scienteer after completing the final SEF competition in which they participated. Those who did were presented with a hyperlink to the SEF survey. Scienteer does not send out reminder emails, and no incentives were offered for remembering to sign back in and participate in the survey.

Since 2016, when we began surveying the national Scienteer cohort of SEF students, 135,000 parents have consented and more than 4,000 students have completed surveys, an overall response rate of about 3%. Given that student participation in the surveys involves an indirect, single electronic invitation without incentive or follow-up, this level of response would be expected [34–36]. Because the students who participate are not personally identifiable, they can share their opinions openly. Also, because these “subjects” are anonymous, we can make the original survey data itself public in supporting information. Other types of research approaches that involve personal student interviews or comparison of student opinions pre vs. post science fair experience would be valuable but are outside the scope of our research.

The survey used for the current study can be found in supporting information (S1 Survey). The current version is similar to the original survey adopted in 2015 for use with combined groups of regional high school students and national bioscience post high school students [37, 38]. Since then, new questions have been added. Major changes included new questions about level of SEF competition, interest in a career in S&E, and whether SEF experience increased S&E interest in 2016/17 [21]; about student ethnicity in 2018-19 [22]; and about location of the student’s high school (urban, suburban, rural) in 2019/20.

Survey data were summarized with frequency counts and percentages. Significance of potential relationships between data items was assessed using Chi-square contingency tables for independent groups. Results are presented both graphically to make overall trends easier to appreciate and in tables to show the actual numbers. A probability value of p = 0.05 or smaller was accepted as statistically significant but actual p values are shown. No adjustments were made for multiple comparisons.

## Results

### Overview of survey responses

Table 1 shows Scienteer registration numbers and ethnicity trends for the three years covered by this report. In 2019/20 and 2021/22 almost 40,000 students registered each year for SEFs. About 40% fewer students registered with Scienteer in 2020/21. In relationship to U.S. high school data (last row of Table 1) [39], obvious differences in ethnicity between Scienteer registration students and the overall U.S. student population were overrepresentation of Asian students and underrepresentation of Black, Hispanic, and White students. Also, compared to U.S. high school data and to survey respondents (below), many more Scienteer students indicated “Other” as reported previously [22].

**Table 1.** Scienteer Registration and Ethnicity

| Data Set | All Students | Asian | Black | Hispanic | White | Other* |
| --- | --- | --- | --- | --- | --- | --- |
|  | survey year | % of students |  |  |  |  |
|  | # of students |  |  |  |  |  |
| Scienceteer SEF registration | 2019/20 | 14.6 | 6.6 | 24.5 | 26.1 | 28.1 |
|  | 38570 | 5365 | 2504 | 9207 | 10010 | 11484 |
|  | 2020/21 | 15.7 | 5.7 | 15.3 | 24.3 | 39.0 |
|  | 23646 | 3720 | 1345 | 3619 | 5746 | 9216 |
|  | 2021/22 | 12.5 | 6.2 | 17.6 | 23.6 | 40.1 |
|  | 39148 | 4908 | 2409 | 6875 | 9246 | 15710 |
|  | Totals – 3 yrs | 14.3 | 6.1 | 19.1 | 24.7 | 35.7 |
|  | 101364 | 13993 | 6258 | 19701 | 25002 | 36410 |
| US High School** | 2019/20 | 5.5 | 14.7 | 27.0 | 48.0 | 4.8 |
|  | 15227541 | 838619 | 2231971 | 4106253 | 7314416 | 736282 |
| *Includes American Indian; Hawaiian Native/Pacific Islander; various mixed-race combinations; and students who choose not to answer. **from NCES Table 216.60 [33] |  |  |  |  |  |  |

Most of the surveys submitted by students (83%) were complete and non-duplicates. In the completed surveys, students answered almost all the questions (>96%) except the question about level of SEF competition, which was less, about 80%. The completed surveys were used for data analyses, and year by year datasets can be found in Supporting Information (S1 Datasets).

In our previous work, we analyzed survey results for two consecutive years to increase consistency, reliability, and reproducibility. However, for the current analysis we utilized three years to allow for a possible effect of COVID in 2020/21. Fig 1 shows year-by-year student survey demographic trends. Consistent with 40% decrease in overall Scienteer registration, almost 40% fewer students completed SEF surveys in 2020/21. Student participation grade, school location (except the percent of students who indicated rural schools), gender, and ethnicity were similar for all three survey years. Supplemental Table 1 (S1 Table) shows few differences in the entire year-by-year set of results for 100+ possible survey questions and answers regarding student demographics, opinions about SEF, help received, obstacles encountered, and ways of overcoming obstacles. Some small differences might have been COVID-related, e.g., more students doing individual projects and fewer getting help from other students in 2020/21. For subsequent analyses, the 2019/20, 2020/21 and 2021/22 student survey results were combined.

**Fig 1.**
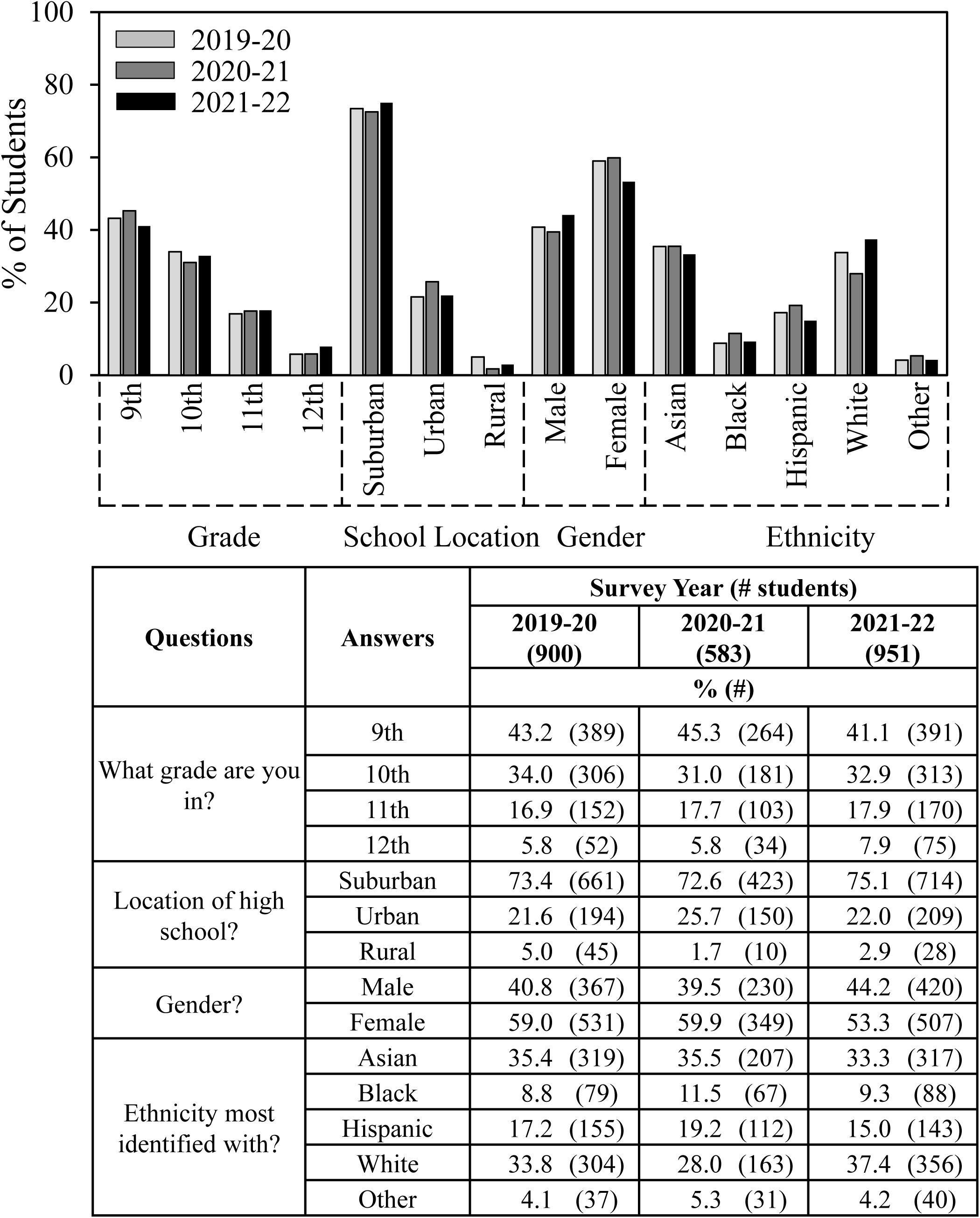
Student survey demographics – year-to-year similarity

### School Location and Student Ethnicity

Fig 2 shows survey trends for student ethnicity and school location. Ethnicity findings for Scienteer students who completed surveys followed a similar pattern overall compared to those who had registered with Scienteer except the category of students who selected “Other” was less than 5% rather than over 30%. Compared to national student enrollment (Table 1), Asian students participating in SEFs were overrepresented almost 6-fold, whereas Black, Hispanic, and White students were underrepresented by about one third.

**Fig 2.**
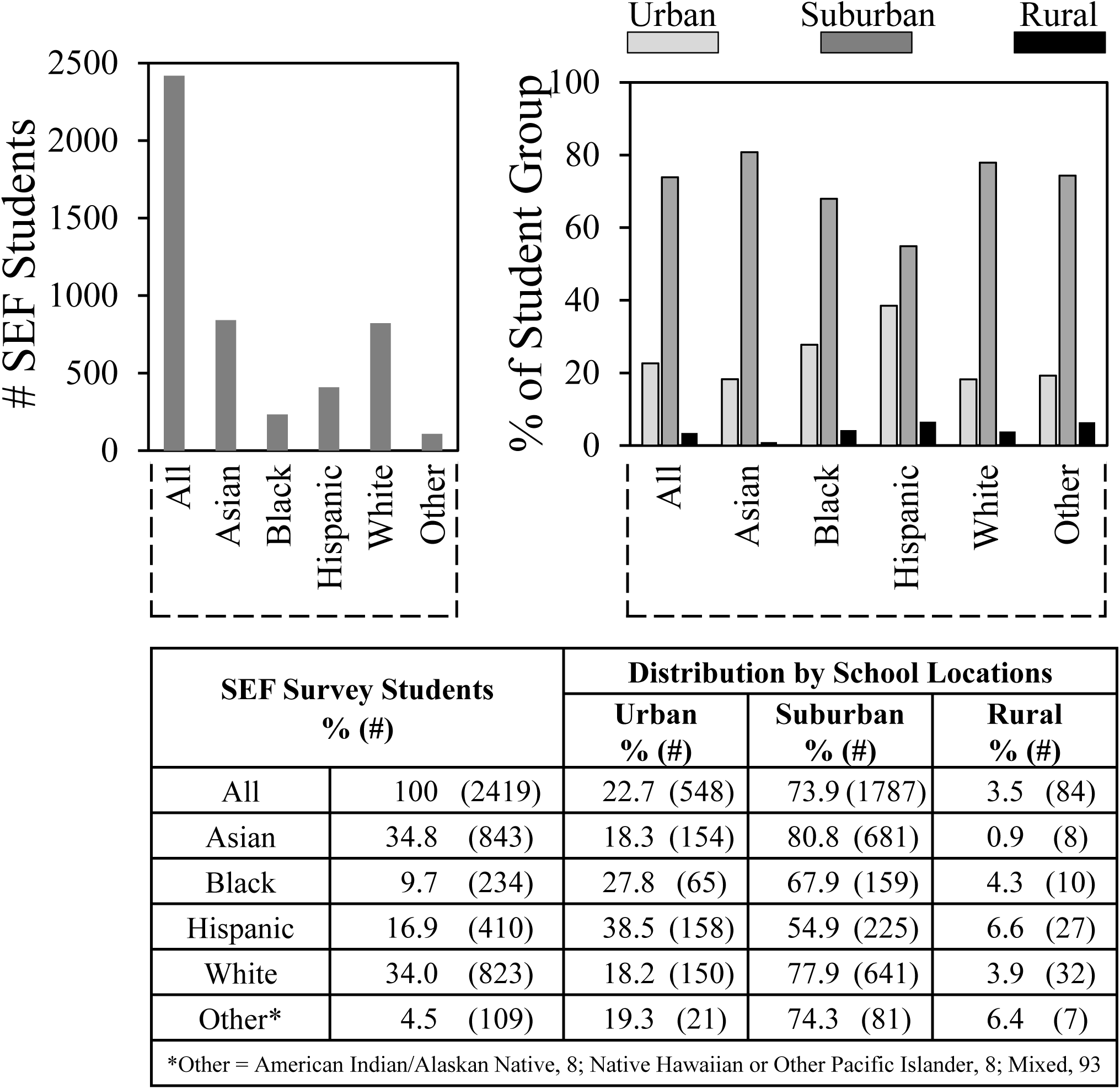
Student ethnicity and school location

Most students indicated that they were from suburban schools – about 80% for Asian and White students, 68% for Black students, and 55% for Hispanic students. National student data shows enrollment of Black and Hispanic students in urban vs. suburban schools at 45.4% and 41.3% respectively [39]. Therefore, representation of urban Hispanic students in SEFs (38.5%) was close to their overall urban representation. Less than 4% of the students indicated that they were from rural schools, which contrasts sharply with student attendance at rural school that ranges from 7% of Asian students to 28% of White students [39].

### School Location, Student Experience and SEF Outcomes

Figs 3-8 compare results regarding SEF outcomes in relationship to school location for students’ overall experiences (Fig 3); ethnicity (Fig 4 & 5); and gender (Fig 6-8). In these figures we only compare students from suburban vs. urban schools because so few students were from rural schools.

**Fig 3.**
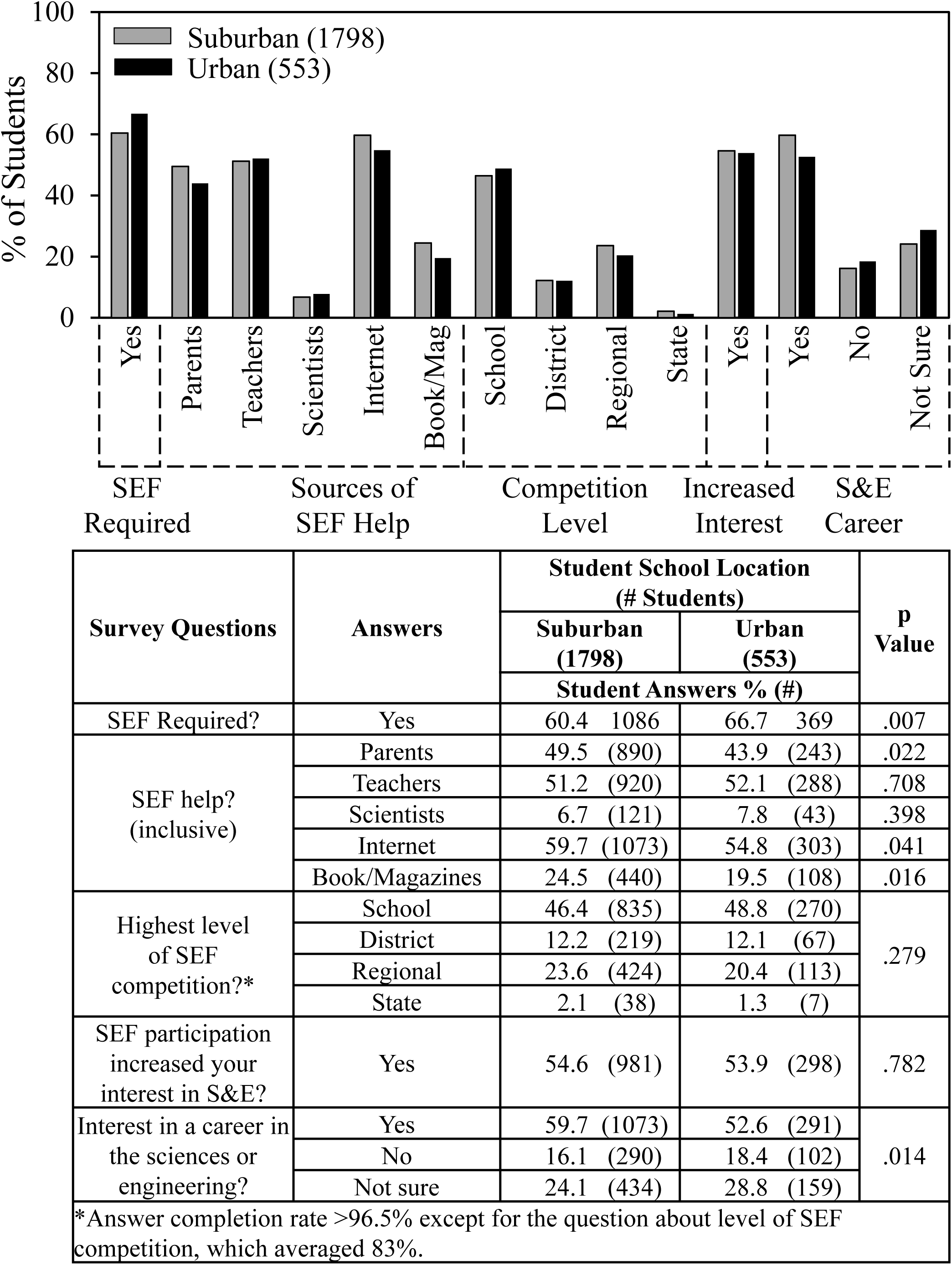
School location and SEF help and outcomes

Fig 3 shows that most students who participated in SEFs were required to do so, 60% from suburban schools and 67% from urban schools. No differences were evident between suburban and urban students in the level of SEF competition or the students’ responses to the question regarding whether SEF participation increased their interests in S&E. About half the students received help from teachers and less than 10% received help from scientists.

Compared to students from urban schools, those from suburban schools were slightly more likely (by a few percentage points) to receive help from parents, use the internet and books and magazines, and indicate an interest in a career in the sciences or engineering.

### School Location, Ethnicity and SEF Experience

Fig 4 (control for Fig 5) confirms and extends previous findings from our recent study on SEF experiences according to student ethnicity [22]. Asian and Hispanic students had more successful SEF experiences than Black and White students according to level of competition reached and on whether students indicated that SEF participation increased their interests in S&E. Asian students were the most likely to get help from scientists and to be interested in a career in the sciences or engineering; Black students were least likely. White students were most likely to get help from parents and help fine tuning their SEF reports.

**Fig 4.**
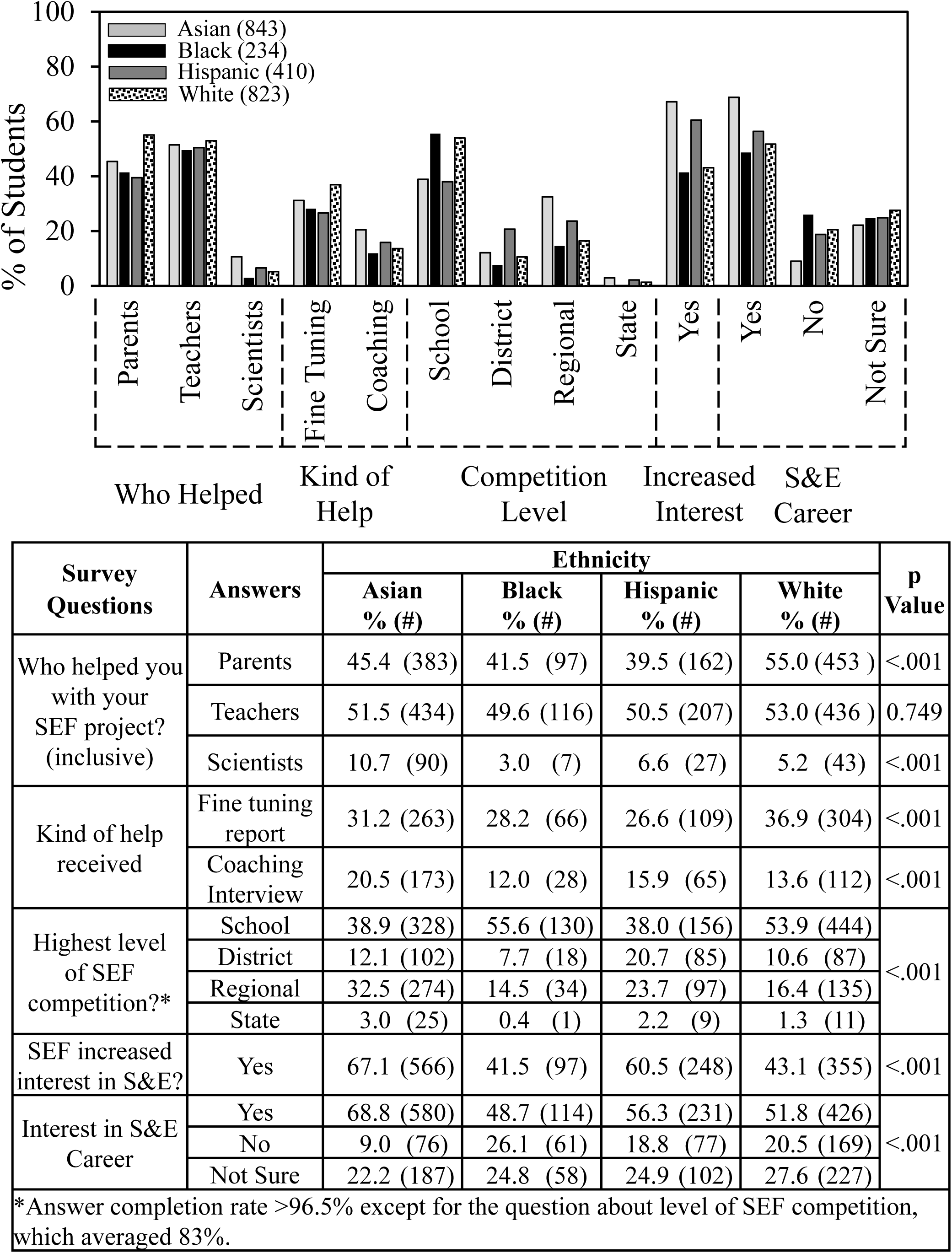
Ethnicity and SEF experience

In Fig 5, we re-ordered the data from Fig 4 and found that students’ experiences by ethnicity were mostly independent of school location. Numerical and statistical details for Fig 5 can be found organized by ethnic group in supplemental data (S2 Table). The only slight differences (by a few percentage points) were that Asian students from suburban vs. urban schools were more likely to advance to SEFs beyond the school level and more likely to indicate an interest in a career in the sciences or engineering. Hispanic students from suburban schools vs. urban were more likely to get help fine tuning their reports. White students from urban vs. suburban schools were more likely to get help from scientists.

**Fig 5.**
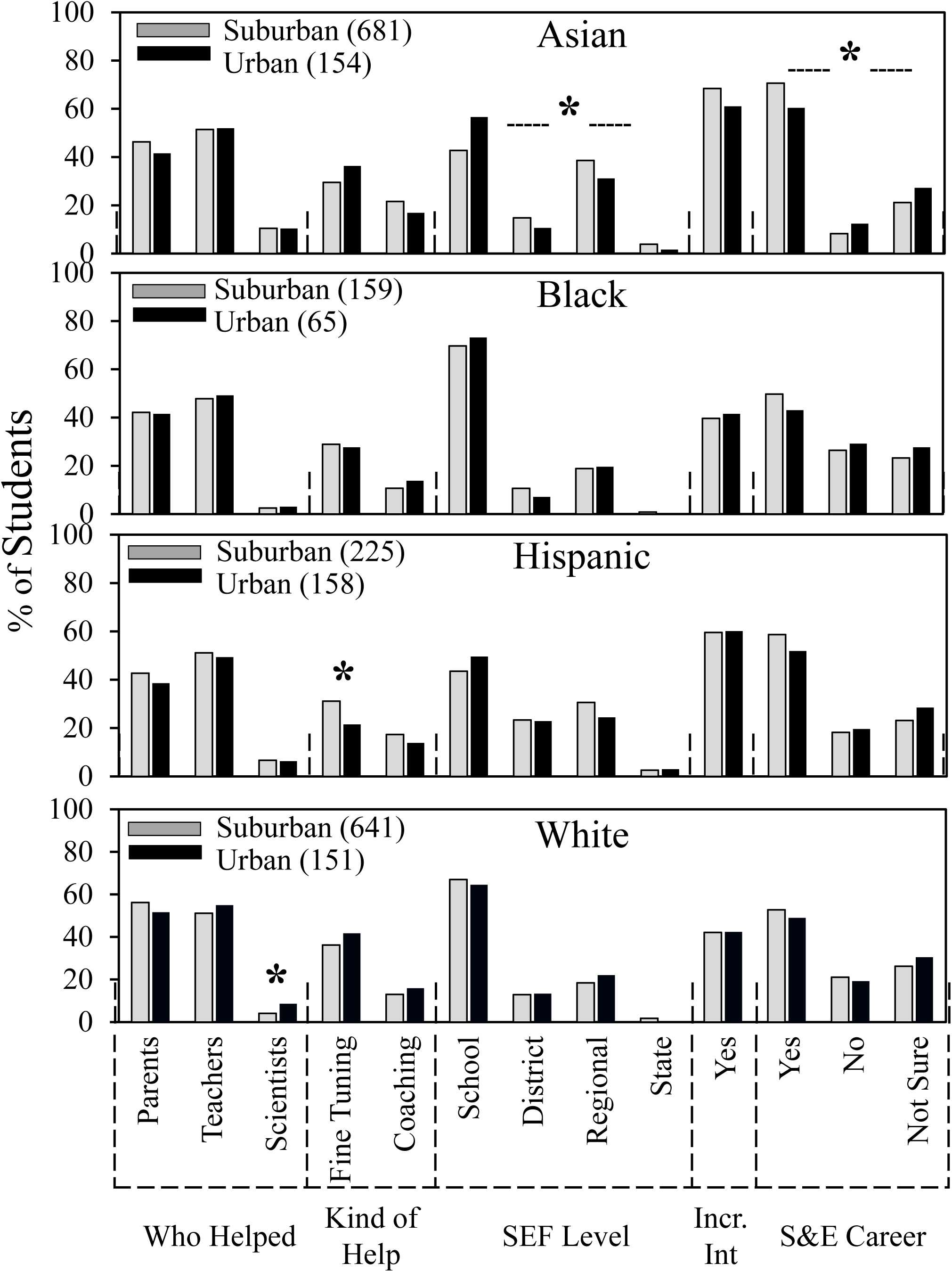
School location, ethnicity, and SEF experience

### School Location, Gender and SEF Experience

Fig 6 shows survey trends for student gender, location, and ethnicity. A majority of students who completed surveys were female (1.37:1). That ratio was lower for suburban schools (1.28:1) and higher for urban schools (1.69:1). In relationship to ethnicity, for Hispanic and White students, females outnumbered male students 1:44:1 and 1.37:1 respectively; For Asian students, the ratio was almost even (1.10:1); and for Black students, females far outnumbered males (2.47:1).

**Fig 6.**
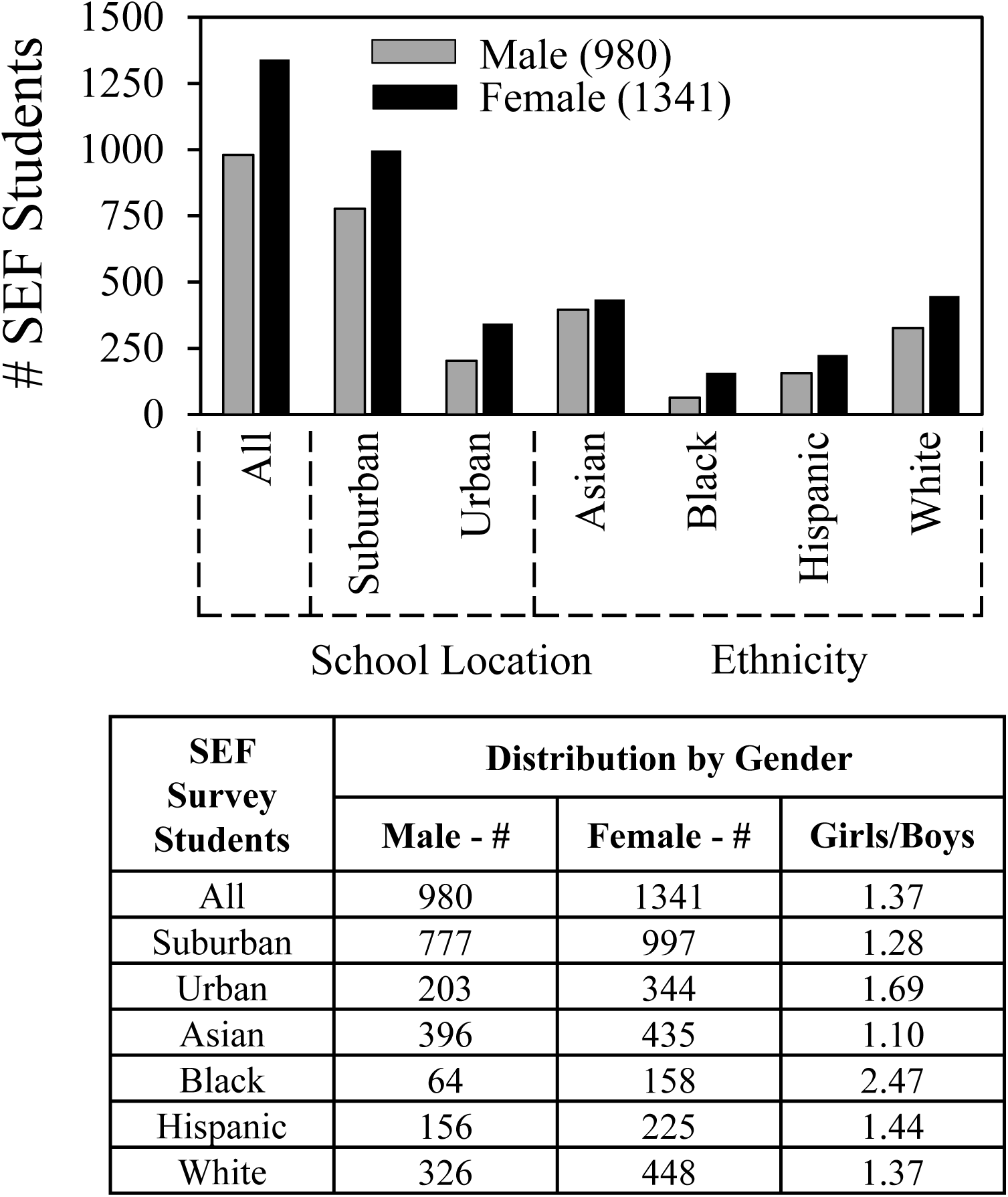
Gender, school location and ethnicity

Fig 7 (control for Fig 8) shows that male and female students received a similar degree of help from parents and teachers and reached similar levels of SEF competitions. Males were more likely by a few percentage points to receive help from scientists, to indicate that SEFs increased their interests in S&E, and to be interested in a S&E career.

**Fig 7.**
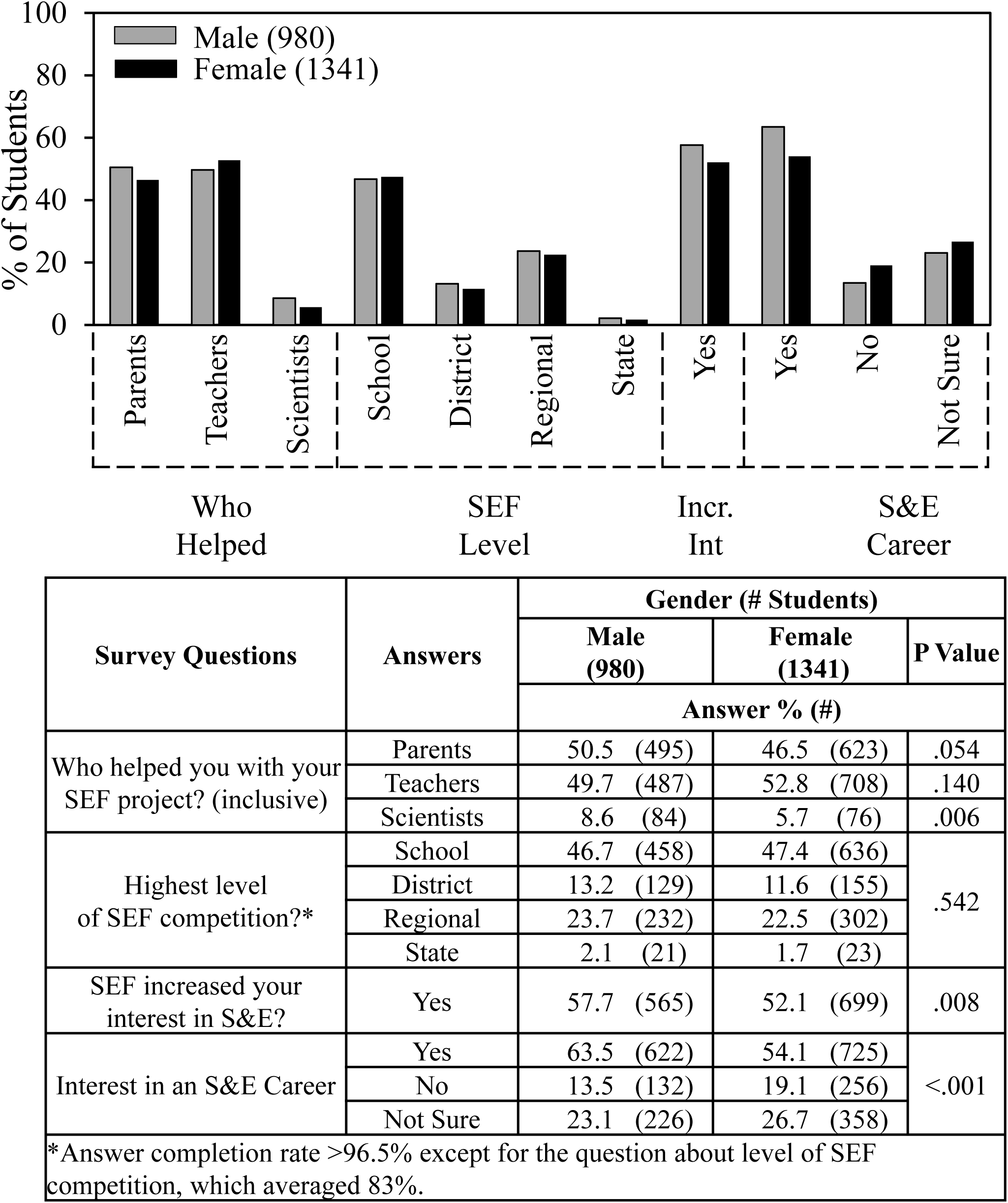
Gender and SEF experience

In Fig 8, we re-ordered the data from Fig 7 and found that students’ experiences by gender were mostly independent of school location. Numerical and statistical details for Fig 6 can be found organized by ethnic group in supplemental data (S3 Table). The only slight difference (by a few percentage points) was that males from suburban schools received more help from parents compared to males from urban schools.

**Fig 8.**
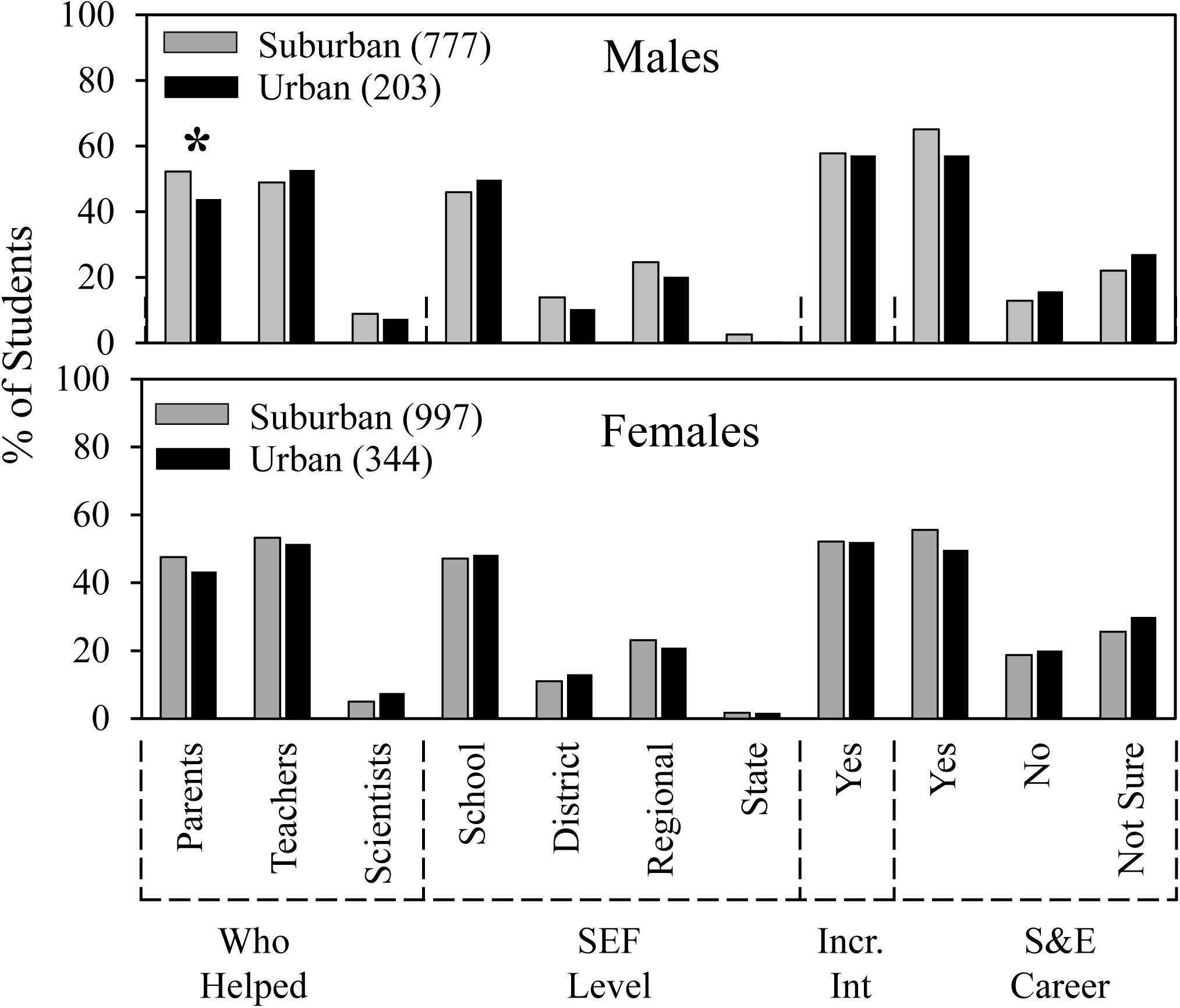
School location, gender, and SEF experience

## Discussion

The overall goal of our research is descriptive. We aim to develop a base of knowledge regarding student experiences in high school SEFs that will help identify best practices leading to more effective, inclusive, and equitable SEF learning opportunities thereby enhancing successful student participation and outcomes such as described elsewhere [40, 41].

In this paper, we report location trends in U.S. student participation in high school science and engineering fairs in relationship to student experiences and SEF outcomes. Beginning in 2019/20, we added a question to our ongoing Scienteer SEF surveys to ask students *Location of high school?* with the possible answers, *urban, suburban, or rural*. About 2500 students who registered and managed SEF projects through the Scienteer website completed surveys over the three school years 2019/20, 2020/21, 2021/22. As will be discussed, although most students participating in SEFs came from suburban schools, students participating in SEFs and coming from urban schools indicated equivalent SEF experiences. Very few students participating in SEFs indicated that they were from rural schools.

During 2020/21, we observed about a 40% decrease in both students who signed up for SEFs using Scienteer and the number of students who filled out surveys. To increase the consistency, reliability, and reproducibility of our findings, we have in the past analyzed survey results for two consecutive years. However, for the current analysis we utilized three years to allow for a decrease in SEF registration in 2020/21 because COVID disrupted many SEFs. Students frequently attended school virtually and were unable to do SEF projects. However, based on their survey answers, those students who did participate in SEFs during 2020/21 had similar experiences compared to the other years with only some small differences that might be COVID-related such as fewer students doing team projects and fewer indicating they received help from other students.

One limitation of our study is that we treat the Scienteer SEF population as a national group. However, it should be recognized that these students may not be truly representative of a national sample since they come from only 7 U.S. states and only attend high schools where SEFs are available. In addition, we cannot be sure that the 3% response rate of survey respondents is representative of the high school student population participating in SEFs. However, the answers of the national cohort of students surveyed through Scienteer regarding opinions about whether SEFs should be required, sources of help, types of help, obstacles encountered, and means of overcoming obstacles closely overlap not only from year to year, but also with most answers of the regional high school students we surveyed initially with surveys distributed directly and the response rate 57% [37, 38].

Consistent with previous findings [22], Asian students were overrepresented in SEF participation compared to the U.S. school population, whereas Black, Hispanic, and White students were underrepresented. Adding the new location data, we learned that for all ethnic groups, the majority of SEF students indicated that they came from suburban schools – Asian, 81%; Black, 68%; Hispanic, 55%; and White, 78%. Although more students from suburban schools participated in SEFs compared to students from urban schools, overall student experiences and outcomes for each ethnic each group were very similar including level of SEF competition and whether SEF participation increased a student’s interest in S&E. While some statistically significant differences (p<.05) were observed between students from suburban and urban schools, all involved only small percentages of students.

Another limitation of our study is that we cannot be sure that the students correctly self-reported their school locations. Some evidence favoring this assumption is that a higher proportion of Black and Hispanic students compared to Asian and White students indicated they were from urban schools as would be expected based on national education data. Indeed, the percentage of Hispanic students in our surveys who indicated that they participated in SEFs coming from urban schools, 38.5%, was similar to the percentage of Hispanic students reported to attend urban schools, 41.3% [39].

Previous research showed that access to outside of school facilities, research resources, and help from scientists enhanced students’ SEF experiences [22, 28, 29]. These findings are consistent with the observation in the education literature that socioeconomic resources are key factors in educational achievement [32, 42, 43]. Although segregation across districts and communities continues [44–46], it also has become clear that economic and other resources of individual schools within a single school district can be just as unbalanced and segregated as across districts [47–49]. Without future research to learn more about the SEF students’ socioeconomic situation, we cannot generalize the findings regarding SEF experience in relationship to school location. However, it seems most likely that opportunities at the school level rather than the district or community level are most important given the observation that Asian and Hispanic students indicated the most positive SEF outcomes, but Asian students had the highest percentage of SEF participants from suburban vs. urban schools – 81% vs. 18%; whereas Hispanic students had the most balanced representation of participants from suburban vs. urban schools – 55% vs. 39%.

Education research typically uses urban, suburban, and rural designations to describe learning environments [50], whereas the U.S. Department of Education uses more nuanced definitions including the category *town* as a location distinct from *city*, *suburban*, and *rural* [51]. According to the latter, 19.5% of students overall including 28% of White students attend rural schools [39]. By contrast, in our surveys only 3.4% -- 84 out of 2419 – indicated that they came from rural schools. Underrepresentation of White students in SEFs might be a consequence of the low participation of students coming from rural schools.

Various reasons could explain the apparent low SEF participation of students coming from rural schools. One is that rural schools are not using Scienteer. Even if there is a rural school or district SEF, the school might not consider entering students in a regional competition requiring Scienteer registration if travel and overnight accommodations with their associated costs would be necessary to compete in a regional fair. We have no information about the locations where SEFs are held. Another factor might be smaller school size. Only 46% of high schools nationally participate in SEFs, and smaller schools are half as likely than larger schools to offer such programs [52]. Organizing a SEF competition requires a critical mass to make the effort worthwhile. At a rural school with small enrollment, it may not be cost effective for the administration or time-effective for the teachers to organize a SEF. Finally, lack of rural SEFs might be another reflection of decreased STEM education opportunities known to exist in rural communities [53–56]. Distinguishing between the foregoing possibilities will require future research.

Previous work by others showed no gender differences in SEF outcomes but clear differences in SEF subject area preference, e.g., life science for females and physical science for males [57–59]. In our surveys, we observed no gender differences in help from parents and teachers and levels of SEF competitions. However, males were more likely by a few percentage points to receive help from scientists, to indicate that SEFs increased their interests in S&E, and to be interested in a S&E career. Nevertheless, none of these gender differences were affected significantly by whether students indicated they came from suburban vs. urban schools.

In conclusion, based on surveys of high school students who participated in SEFs over the past three years, we have learned three new and important features about their experiences. First, most students participating in SEFs indicated that they come from suburban schools. Second, students participating in SEFs and coming from urban schools have equivalent SEF experiences to those from suburban schools. And third, very few students participating in SEFs apparently come from rural schools. In addition, the new findings confirm the previous observation that overall, Asian and Hispanic students indicate better high school SEF experiences than Black and White students.

## Acknowledgments

We are grateful to Russell Cowen and Rocky Slavin, managers of Scienteer Technologies, who incorporated the parental consent and SEF survey REDCap links into the Scienteer website and continue to provide ongoing oversight and management of survey access. Drs. Elise Christopher from the National Center for Education Statistics and Maya Riser-Kositsky from *Education Week* advised us about using U.S. Department of Education National Center for Education Statistics.

## Supporting information

S1 Data set. Excel dataset showing all of the survey questions and answers.

S1 Survey. Survey questions.

S1 Table. Comparison of year-to-year survey responses.

S2 Table. School location, ethnicity, and students’ SEF experiences.

S3 Table. School location, gender, and students’ SEF experiences.

---

**Supplemental Table 1.**
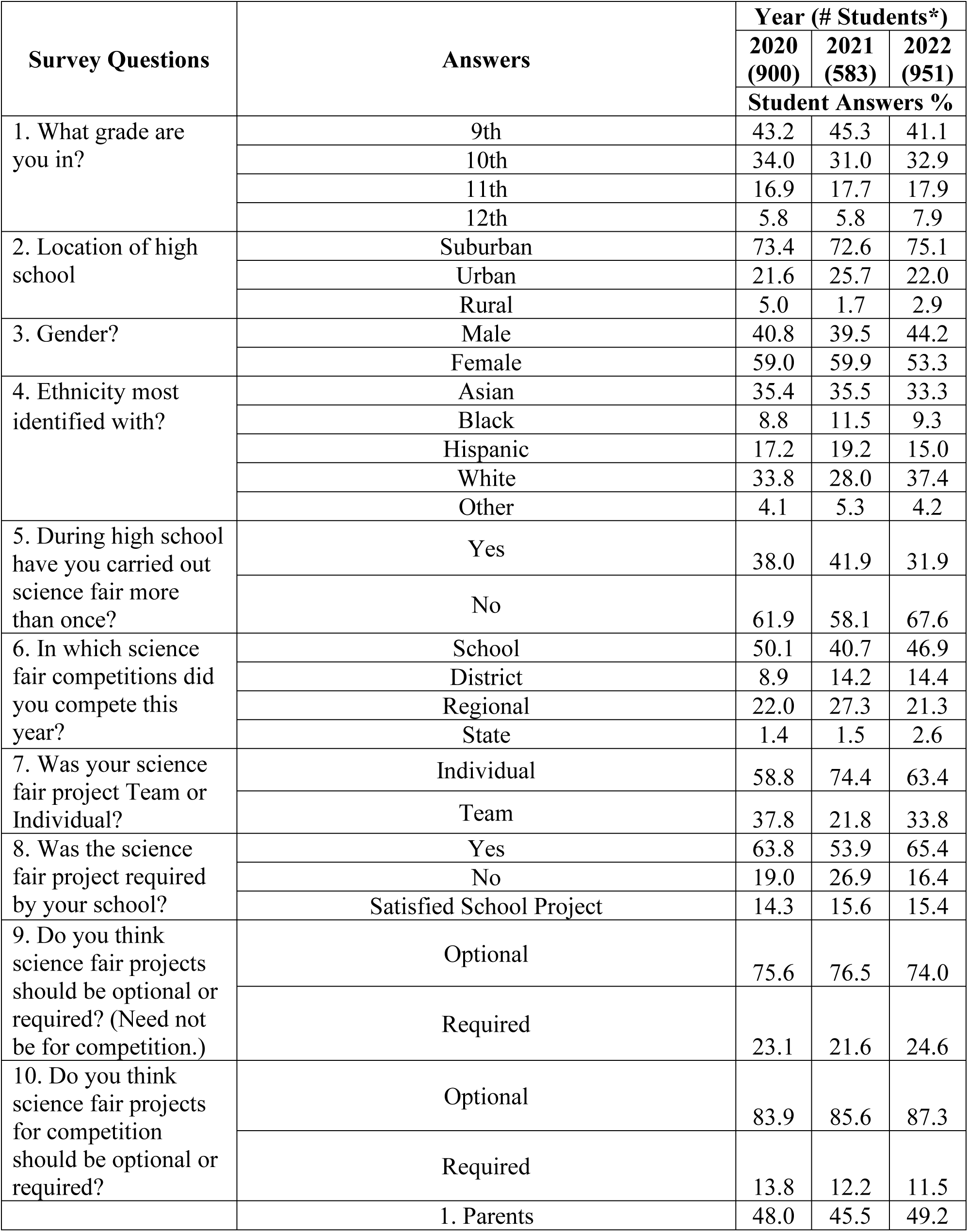
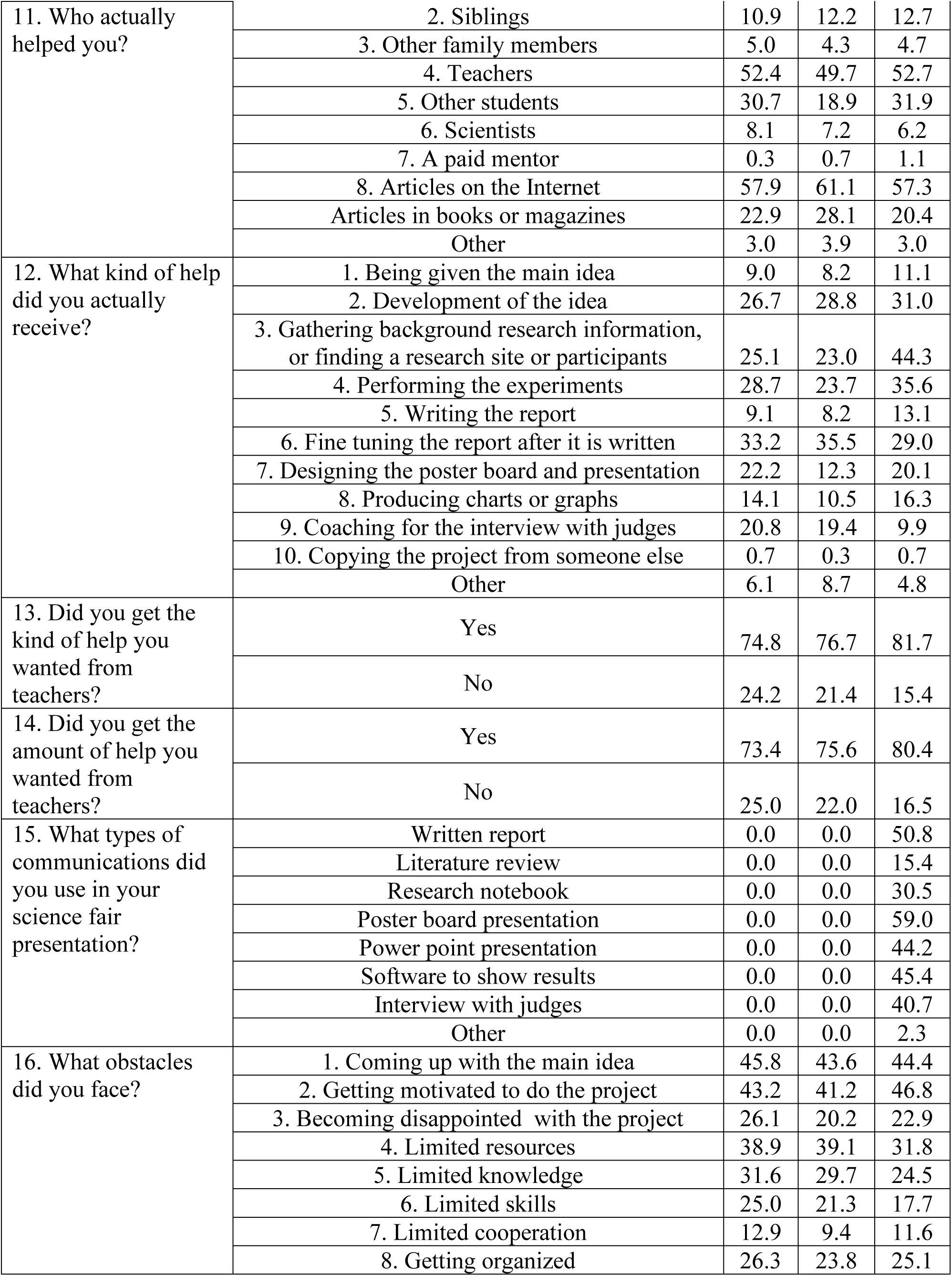
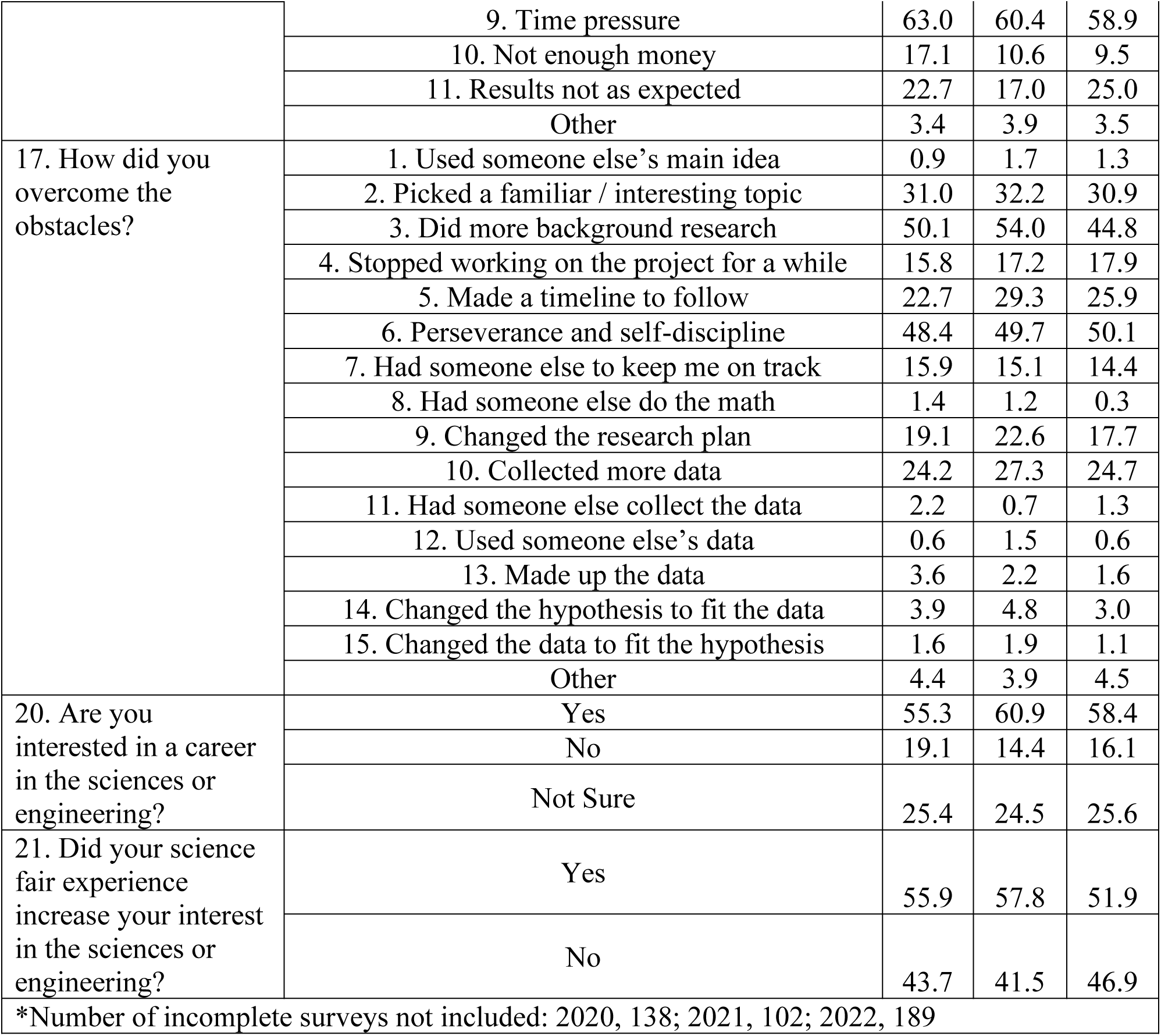
Student survey answers year-to-year

**Supplemental Table 2.** School location, ethnicity, and SEF experience

| Survey Questions | Answers | School Location |  | p Value |
| --- | --- | --- | --- | --- |
|  |  | Suburban | Urban |  |
|  |  | Student Answers % (#) |  |  |
| Asian Students # |  | 681 | 154 |  |
| Who helped you with your SEF project?<br>(more than one answer possible) | Parents | 46.3 (315) | 41.6 (64) | 0.290 |
|  | Teachers | 51.4 (350) | 51.9 (80) | 0.901 |
|  | Scientists | 10.4 (71) | 10.4 (16) | 0.989 |
| Kind of help received | Fine tuning report | 29.5 (201) | 36.4 (56) | 0.096 |
|  | Coaching interview | 21.6 (147) | 16.9 (26) | 0.193 |
| Highest level<br>of SEF competition? | School | 42.8 (257) | 56.6 (69) | 0.038 |
|  | District | 14.8 (89) | 10.7 (13) |  |
|  | Regional | 38.6 (232) | 31.1 (38) |  |
|  | State | 3.8 (23) | 1.6 (2) |  |
| SEF Increased Interest | Yes | 68.4 (466) | 61.0 (94) | 0.078 |
| Interest in S&E Career | Yes | 70.6 (481) | 60.4 (93) | 0.041 |
|  | No | 8.2 (56) | 12.3 (19) |  |
|  | Not Sure | 21.1 (144) | 27.3 (42) |  |
| Black Students # |  | 159 | 65 |  |
| Who helped you with your SEF project?<br>(more than one answer possible) | Parents | 42.1 (67) | 41.5 (27) | 0.934 |
|  | Teachers | 47.8 (76) | 49.2 (32) | 0.846 |
|  | Scientists | 2.5 (4) | 3.1 (2) | 0.813 |
| Kind of help received | Fine tuning report | 28.9 (46) | 27.7 (18) | 0.852 |
|  | Coaching interview | 10.7 (17) | 13.8 (9) | 0.504 |
| Highest level<br>of SEF competition? | School | 69.7 (85) | 73.2 (41) | 0.831 |
|  | District | 10.7 (13) | 7.1 (4) |  |
|  | Regional | 18.9 (23) | 19.6 (11) |  |
|  | State | 0.8 (1) | 0.0 (0) |  |
| SEF Increased Interest | Yes | 39.6 (63) | 41.5 (27) | 0.791 |
| Interest in S&E Career | Yes | 49.7 (79) | 43.1 (28) | 0.632 |
|  | No | 26.4 (42) | 29.2 (19) |  |
|  | Not Sure | 23.3 (37) | 27.7 (18) |  |
| Hispanic Students # |  | 225 | 158 |  |
| Who helped you with your SEF project?<br>(more than one answer possible) | Parents | 42.7 (96) | 38.6 (61) | 0.426 |
|  | Teachers | 51.1 (115) | 49.4 (78) | 0.737 |
|  | Scientists | 6.7 (15) | 6.3 (10) | 0.895 |
| Kind of help received | Fine tuning report | 31.1 (70) | 21.5 (34) | 0.038 |
|  | Coaching interview | 17.3 (39) | 13.9 (22) | 0.369 |
| Highest level<br>of SEF competition? | School | 43.5 (84) | 49.6 (65) | 0.627 |
|  | District | 23.3 (45) | 22.9 (30) |  |
|  | Regional | 30.6 (59) | 24.4 (32) |  |
|  | State | 2.6 (5) | 3.1 (4) |  |
| SEF Increased Interest | Yes | 59.6 (134) | 60.1 (95) | 0.911 |
| Interest in S&E Career | Yes | 58.7 (132) | 51.9 (82) | 0.384 |
|  | No | 18.2 (41) | 19.6 (31) |  |
|  | Not Sure | 23.1 (52) | 28.5 (45) |  |
| **White Students #** |  | **641** | **151** |  |
| Who helped you with your SEF project?<br>(more than one answer possible) | Parents | 56.2 (360) | 51.7 (78) | 0.316 |
|  | Teachers | 51.2 (328) | 55.0 (83) | 0.401 |
|  | Scientists | 4.1 (26) | 8.6 (13) | 0.02 |
| Kind of help received | Fine tuning report | 36.2 (232) | 41.7 (63) | 0.206 |
|  | Coaching interview | 12.9 (83) | 15.9 (24) | 0.341 |
| Highest level<br>of SEF competition? | School | 67.0 (349) | 64.6 (82) | 0.709 |
|  | District | 12.9 (67) | 13.4 (17) |  |
|  | Regional | 18.4 (96) | 22.0 (28) |  |
|  | State | 1.7 (9) | 0.0 (0) |  |
| SEF Increased Interest | Yes | 42.1 (270) | 42.4 (64) | 0.953 |
| Interest in S&E Career | Yes | 52.7 (338) | 49.0 (74) | 0.512 |
|  | No | 21.1 (135) | 19.2 (29) |  |
|  | Not Sure | 26.2 (168) | 30.5 (46) |  |

**Supplemental Table 3.** School location, gender, and SEF experience

| Survey Questions | Answers | Location |  | p Value |
| --- | --- | --- | --- | --- |
|  |  | Suburban | Urban |  |
|  |  | Student Answers % (#) |  |  |
| Males # |  | 777 | 203 |  |
| Who helped you with your SEF project? (inclusive) | Parents | 52.3 (406) | 43.8 (89) | 0.033 |
|  | Teachers | 48.9 (380) | 52.7 (107) | 0.334 |
|  | Scientists | 8.9 (69) | 7.4 (15) | 0.499 |
| Highest level of SEF competition? | School | 45.9 (357) | 49.8 (101) | 0.105 |
|  | District | 13.9 (108) | 10.3 (21) |  |
|  | Regional | 24.6 (191) | 20.2 (41) |  |
|  | State | 2.6 (20) | 0.5 (1) |  |
| SEF increased your interest in S&E | Yes | 57.8 (449) | 57.1 (116) | 0.869 |
| Interest in an S&E Career | Yes | 65.1 (506) | 57.1 (116) | 0.110 |
|  | No | 12.9 (100) | 15.8 (32) |  |
|  | Not Sure | 22.0 (171) | 27.1 (55) |  |
| Females # |  | 997 | 344 |  |
| Who helped you with your SEF project? (inclusive) | Parents | 47.5 (474) | 43.3 (149) | 0.175 |
|  | Teachers | 53.3 (531) | 51.5 (177) | 0.563 |
|  | Scientists | 5.0 (50) | 7.6 (26) | 0.079 |
| Highest level of SEF competition? | School | 47.1 (470) | 48.3 (166) | 0.688 |
|  | District | 11.0 (110) | 13.1 (45) |  |
|  | Regional | 23.1 (230) | 20.9 (72) |  |
|  | State | 1.7 (17) | 1.7 (6) |  |
| SEF increased your interest in S&E | Yes | 52.2 (520) | 52.0 (179) | 0.969 |
| Interest in an S&E Career | Yes | 55.6 (554) | 49.7 (171) | 0.159 |
|  | No | 18.8 (187) | 20.1 (69) |  |
|  | Not Sure | 25.6 (255) | 29.9 (103) |  |

## References Cited

1. Furtak EM, Penuel WR. Coming to terms: Addressing the persistence of “hands-on” and other reform terminology in the era of science as practice. Science education. 2019;103(1):167–86.

2. Lederman NG. Contextualizing the relationship between nature of scientific knowledge and scientific inquiry. Science & Education. 2019;28(3):249–67.

3. Osborne J. Teaching scientific practices: Meeting the challenge of change. Journal of Science Teacher Education. 2014;25(2):177–96.

4. National Research Council. A Framework for K-12 Science Education: Practices, Crosscutting Concepts, and Core Ideas. Washington, D.C.: National Academies Press; 2012.

5. NGSS Lead States. Next Generation Science Standards For States, By States. Volume 1: The Standards—Arranged by Disciplinary Core Ideas and by Topics. Washington, D.C.: National Academies Press; 2013.

6. Kook JF, DeLisi J, Fields ET, Levy AJ. Approaches for conducting middle school science fairs: a landscape study. Science Educator. 2020;27(2):71–80.

7. Burgin SR, Sadler TD. Learning nature of science concepts through a research apprenticeship program: A comparative study of three approaches. Journal of Research in Science Teaching. 2016;53(1):31–59.

8. Houseal AK, Abd-El-Khalick F, Destefano L. Impact of a student–teacher–scientist partnership on students’ and teachers’ content knowledge, attitudes toward science, and pedagogical practices. Journal of Research in Science Teaching. 2014;51(1):84–115.

9. Andersson J, Schaben C, Buhs E, Grandgenett N. Phenomenology of Secondary Students’ Experiences in Out-of-School Time Science Research. Science Educator. 2021;28(1):30–9.

10. Rodenbusch SE, Hernandez PR, Simmons SL, Dolan EL. Early engagement in course-based research increases graduation rates and completion of science, engineering, and mathematics degrees. CBE—Life Sciences Education. 2016;15(2):ar20.

11. Hanauer DI, Graham MJ, Sea-Phages, Betancur L, Bobrownicki A, Cresawn SG, et al. An inclusive Research Education Community (iREC): Impact of the SEA-PHAGES program on research outcomes and student learning. Proceedings of the National Academy of Sciences. 2017;114(51):13531–6.

12. Beier ME, Kim MH, Saterbak A, Leautaud V, Bishnoi S, Gilberto JM. The effect of authentic project-based learning on attitudes and career aspirations in STEM. Journal of Research in Science Teaching. 2019;56(1):3–23.

13. U.S. Department of Education National Center for Education Statistics. X2RACE from High School Longitudinal Study of 2009 (HSLS:09) First Follow-up 2013. Available from: https://nces.ed.gov/pubs2014/2014360.pdf.

14. National Academies of Sciences E, and Medicine. Science and engineering for grades 6-12: Investigation and design at the center: National Academies Press; 2019.

15. U.S. Department of Education National Center for Education Statistics. Private Elementary/Secondary School Universe Survey,” 2019-20 2021. Available from: https://nces.ed.gov/programs/digest/d21/tables/dt21_205.10.asp.

16. Miller K, Sonnert G, Sadler P. The influence of students’ participation in STEM competitions on their interest in STEM careers. International Journal of Science Education, Part B. 2018;8(2):95–114.

17. DeLisi J, Kook JF, Levy AJ, Fields E, Winfield L. An examination of the features of science fairs that support students’ understandings of science and engineering practices. Journal of Research in Science Teaching. 2021;58(4):491–519.

18. Fields E, DeLisi J, Kook J, Winfield L, Levy AJ. Parent Involvement in the Science Fair: Helping Students or Hindering Equity? School Community Journal. 2022;32(2).

19. Dabney KP, Tai RH, Almarode JT, Miller-Friedmann JL, Sonnert G, Sadler PM, et al. Out-of-school time science activities and their association with career interest in STEM. International Journal of Science Education, Part B. 2012;2(1):63–79.

20. Grinnell F, Dalley S, Reisch J. High school science fair: Experiences of two groups of undergraduate bioscience students. PloS one. 2021;16(6):e0252627.

21. Grinnell F, Dalley S, Reisch J. High school science fair: Positive and negative outcomes. PLOS ONE. 2020;15(2):e0229237.

22. Grinnell F, Dalley S, Reisch J. High school science fair: Ethnicity trends in student participation and experience. PloS one. 2022;17(3):e0264861.

23. Mernoff B, Aldous AR, Wasio NA, Kritzer JA, Sykes ECH, O’Hagan K. A Reverse Science Fair that Connects High School Students with University Researchers. Journal of Chemical Education. 2017;94(2):171–6.

24. Lakin JM, Ewald ML, Hardy EE, Cobine PA, Marino JG, Landers AL, et al. Getting Everyone to the Fair: Supporting Teachers in Broadening Participation in Science and Engineering Fairs. Journal of Science Education and Technology. 2021:1–20.

25. Koomen MH, Hedenstrom MN, Moran MK. Rubbing elbows with them: Building capacity in STEM through science and engineering fairs. Science Education. 2021;105(3):541–79.

26. Stray S, Gordy X, Sullivan D, Bender S, Cook C, McKone K, et al. Base Pair: 28 years of Sustained High School Biomedical Research Mentorship Driving Health Sciences Career Progression. Journal of STEM Outreach. 2020;3(3):1–10.

27. Todd C. Collaborations between Under-Resourced High School Students and STEM Professionals to Increase Participation in Science and Engineering Fairs. European Journal of Education and Pedagogy. 2022;3(1):1–6.

28. Gifford VD, Wiygul SM. The effect of the use of outside facilities and resources on success in secondary school science fairs. School Science and Mathematics. 1992;92(3):116–9.

29. Bencze JL, Bowen GM. A national science fair: Exhibiting support for the knowledge economy. International Journal of Science Education. 2009;31(18):2459–83.

30. Dalton B, Ingels SJ, Fritch L. High School Longitudinal Study of 2009 (HSLS: 09). 2013 Update and High School Transcript Study: A First Look at Fall 2009 Ninth-Graders in 2013 NCES 2015-037rev. 2016;Table 12.

31. National Science Board. Science & engineering indicators: National Science Board; 2018.

32. Reardon SF, Kalogrides D, Shores K. The geography of racial/ethnic test score gaps. American Journal of Sociology. 2019;124(4):1164–221.

33. Harris PA, Taylor R, Thielke R, Payne J, Gonzalez N, Conde JG. Research electronic data capture (REDCap)—a metadata-driven methodology and workflow process for providing translational research informatics support. Journal of biomedical informatics. 2009;42(2):377–81.

34. Kent R, Brandal H. Improving email response in a permission marketing context. International Journal of Market Research. 2003;45(4):1–13.

35. Van Mol C. Improving web survey efficiency: the impact of an extra reminder and reminder content on web survey response. International Journal of Social Research Methodology. 2017;20(4):317–27.

36. Shih T-H, Fan X. Comparing response rates from web and mail surveys: A meta-analysis. Field methods. 2008;20(3):249–71.

37. Grinnell F, Dalley S, Shepherd K, Reisch J. High school science fair and research integrity. PLOS ONE. 2017;12(3):e0174252.

38. Grinnell F, Dalley S, Shepherd K, Reisch J. High school science fair: Student opinions regarding whether participation should be required or optional and why. PLOS ONE. 2018;13(8):e0202320.

39. U.S. Department of Education National Center for Education Statistics. Public Elementary/Secondary School Universe Survey,” 2019-20 2021. Available from: https://nces.ed.gov/programs/digest/d21/tables/dt21_216.60.asp.

40. Grinnell F. Reinventing Science Fairs. Issues in Science and Technology. 2020;36(3):23–5.

41. Grinnell F, Dalley S. High school science and engineering fairs: Lessons learned. Connected Science Learning 2022;4 (5):https://www.nsta.org/connected-science-learning/connected-science-learning-september-october-2022/high-school-science.

42. Owens A. Income segregation between school districts and inequality in students’ achievement. Sociology of Education. 2018;91(1):1–27.

43. Rauscher E, Shen Y. Variation in the Relationship between School Spending and Achievement: Progressive Spending Is Efficient. American Journal of Sociology. 2022;128(1):189–223.

44. Massey DS, Tannen J. Suburbanization and segregation in the United States: 1970–2010. Ethnic and racial studies. 2018;41(9):1594–611.

45. Fahle EM, Reardon SF, Kalogrides D, Weathers ES, Jang H. Racial segregation and school poverty in the United States, 1999–2016. Race and Social Problems. 2020;12:42–56.

46. Atteberry A, Bischoff K, Owens A. Identifying progress toward ethnoracial achievement equity across US school districts: A new approach. Journal of Research on Educational Effectiveness. 2021;14(2):410–41.

47. Owens A, Reardon SF, Jencks C. Income segregation between schools and school districts. American Educational Research Journal. 2016;53(4):1159–97.

48. Bischoff K, Tach L. The racial composition of neighborhoods and local schools: The role of diversity, inequality, and school choice. City & Community. 2018;17(3):675–701.

49. Schachner JN. Racial Stratification and School Segregation in the Suburbs: Evidence from Los Angeles County. Social Forces. 2022;101(1):309–40.

50. Bradley F, Feldman A. The problematic use of urban, suburban, and rural in science education. Cultural Studies of Science Education. 2021;16(4):1289–313.

51. U.S. Department of Education National Center for Education Statistics. School Locale Definitions 2006. Available from: https://nces.ed.gov/surveys/urbaned/definitions.asp.

52. Banilower ER, Smith PS, Malzahn KA, Plumley CL, Gordon EM, Hayes ML. Report of the 2018 NSSME+. Horizon Research, Inc. 2018.

53. ACT (American College Testing). STEM education in the US: Where we are and what we can do. ACT, Inc. https://www.act.org/content/act/en/research/reports/act …; 2017.

54. Darrah M, Humbert R, Howley C. Differentiating Rural Locale Factors Related to Students Choosing and Persisting in STEM. Research in Higher Education Journal. 2022;42.

55. Lakin JM, Stambaugh T, Ihrig LM, Mahatmya D, Assouline SG. Nurturing STEM talent in rural setting. Phi Delta Kappan. 2021;103(4):24–30.

56. Saw GK, Agger CA. STEM pathways of rural and small-town students: Opportunities to learn, aspirations, preparation, and college enrollment. Educational Researcher. 2021;50(9):595–606.

57. Greenfield TA. An exploration of gender participation patterns in science competitions. Journal of Research in Science Teaching. 1995;32(7):735–48.

58. Sonnert G, Sadler P, Michaels M. Gender aspects of participation, support, and success in a state science fair. School Science and Mathematics. 2013;113(3):135–43.

59. Steegh AM, Höffler TN, Keller MM, Parchmann I. Gender differences in mathematics and science competitions: A systematic review. Journal of Research in Science Teaching. 2019.

